## Supporting information for "Cell type-specific weighting-factors to solve solid organs-specific limitations of single cell RNA-sequencing"

Kengo Tejima *et al.*

**This PDF file includes:**

S1, S2 Figures
S1 to S14 Tables

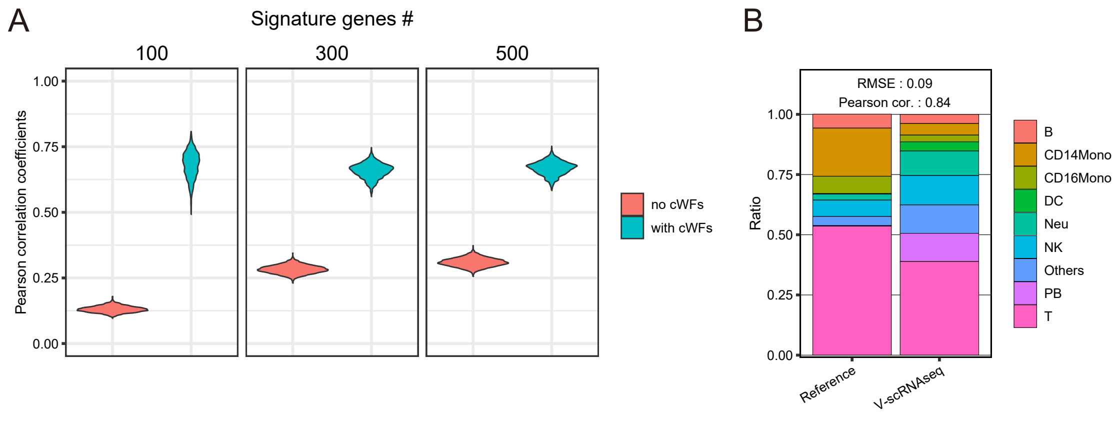


**S1 Fig. Reconstitution and deconvolution of the bulk human PBMC RNA-seq by the composite scRNA-seq.** **(A)** Reconstitution results with and without (no) cWFs are compared. The results with the 100, 300, 500 signature genes are shown. The similarity is shown as violin plot of the Pearson correlation coefficients. The corresponding raw data are found in S6 and S7 Tables. **(B)** Bar graph showing the cell type-ratios computed by the deconvolution method (V-scRNAseq) for each organ. The deconvolution was performed with the cWFs computed using the optimal number of the signature genes for each organ (indicated in the accompanying table S8). The bar graphs are composed of the cell-types computed to be present for each organ by our method. The similarity scores (RMSE: Root Mean Squared Errors, Pearson correlation coefficient) are indicated at the top of the bar. The corresponding raw data of S1 Fig. are found in S6, S7, and S8 Tables.

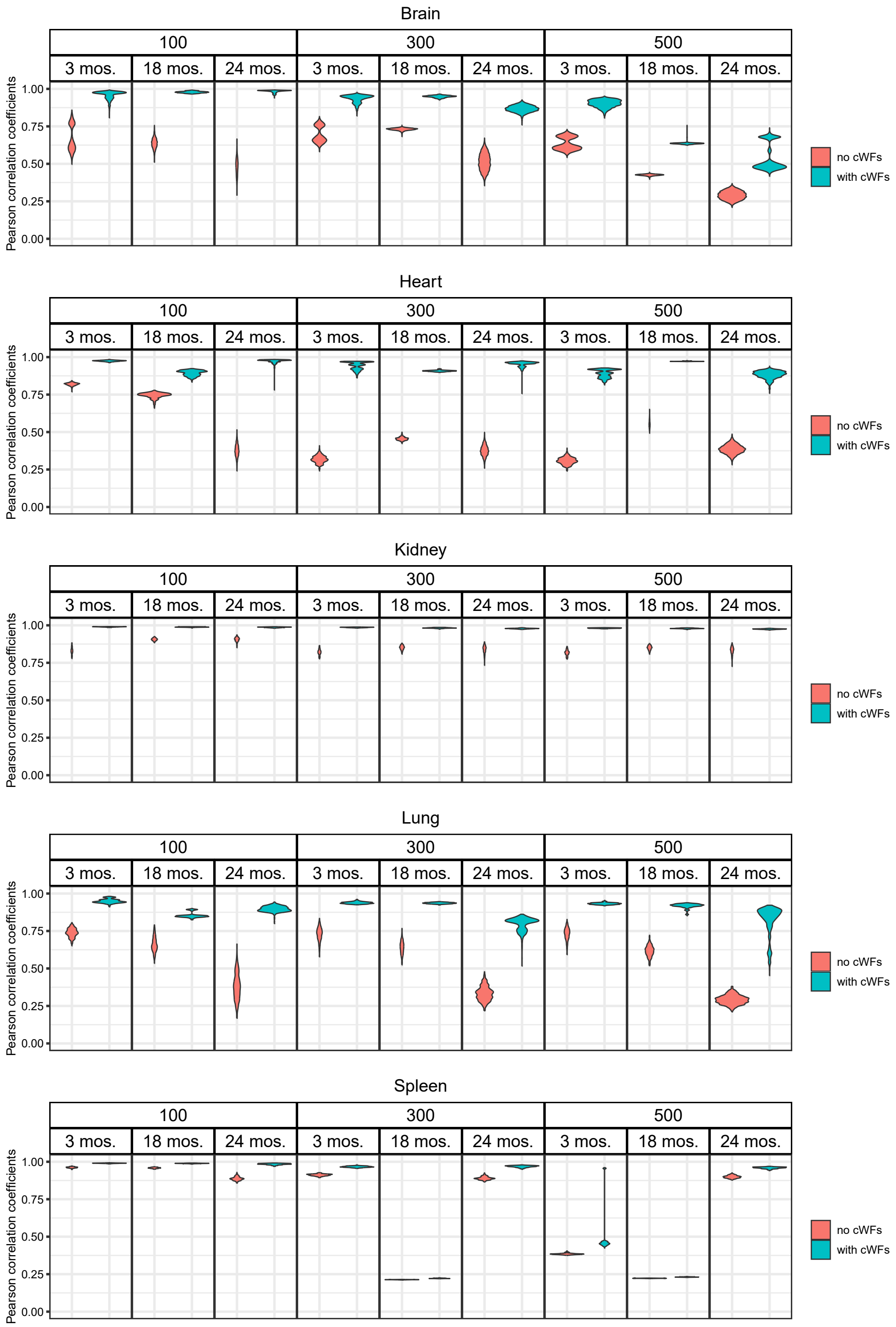


**S2 Fig. Reconstitution of the whole organ RNA-seq of the aging model mouse by the composite scRNA-seq.** The results with and without (no) cWFs are compared for each aging-stage (3 mos., 18 mos. 24 mos.) for each number of the signature gens (100, 300, 500) and for each organ (Brain, Heart, Kidney, Lung, Spleen). The similarity is shown as violin plots of the Pearson correlation coefficients. The corresponding raw data are found in S10 (no cWFs) and S11 (with cWFs) Tables. The corresponding raw data of S2 Fig. are found in S10 and S11 Tables.

All supplementary tables below are available at https://github.com/k-teji/VscRNAseq.

**S1 Table (Separate file)**
The raw data of Fig. 1.

**S2 Table (Separate file)**
The raw data of Fig. 3.

**S3 Table (Separate file)**
The raw data of Fig. 4.

**S4 Table (Separate file)**
The raw data of Fig. 5, B and C.

**S5 Table (Separate file)**
The raw data of Fig. 6.

**S6 Table (Separate file)**
The raw data of S1 Fig. (Pearson correlation coefficients for the result with no cWFs).

**S7 Table (Separate file)**
The raw data of S1A Fig. (Pearson correlation coefficients for the result with cWFs).

**S8 Table (Separate file)**
The raw data of S1B Fig. The best performing result (the lowest RMSE and the highest Pearson correlation coefficient) shown in S1B Fig. is highlighted in light green.

**S9 Table (Separate file)**
The raw data of Fig. 7.

**S10 Table (Separate file)**
The raw data of S2 Fig. (Pearson correlation coefficients for the result with no cWFs).

**S11 Table (Separate file)**
The raw data of S2 Fig. (Pearson correlation coefficients for the result with cWFs).

**S12 Table (Separate file)**
Deconvolution of the aging-mouse whole-organ RNA-seq data. The results omitted due to the lack of the significant improvement by the cWFs in the reconstitution are highlighted in pink. The results used in the analyses shown in Figs. 7, 8, and 9 (the pairwise comparisons between the two aging-stages show $p < 0.01$ by U-test) are highlighted in light-green.

**S13 Table (Separate file)**
The raw data of Fig. 8.

**S14 Table (Separate file)**

The raw data of Fig. 9 (sheet 1) & Fig. 10 (sheet 2).
